# *Abcc6* null mice a model for mineralization disorder PXE show vertebral osteopenia without enhanced intervertebral disc calcification with aging

**DOI:** 10.1101/2021.11.17.468924

**Authors:** Paige K. Boneski, Vedavathi Madhu, Ryan E. Tomlinson, Irving M. Shapiro, Koen van de Wetering, Makarand V. Risbud

## Abstract

Chronic low back pain is a highly prevalent health condition intricately linked to intervertebral disc degeneration. One of the prominent features of disc degeneration that is commonly observed with aging is dystrophic calcification. ATP-binding cassette sub-family C member 6 (ABCC6), a presumed ATP efflux transporter, is a key regulator of systemic levels of the mineralization inhibitor pyrophosphate (PPi). Mutations in ABCC6 result in pseudoxanthoma elasticum (PXE), a progressive human metabolic disorder characterized by mineralization of the skin and elastic tissues. The implications of ABCC6 loss-of-function on pathological mineralization of structures in the spine, however, are unknown. Using the ABCC6^-/-^ mouse model of PXE, we investigated age-dependent changes in the vertebral bone and intervertebral disc. ABCC6^-/-^ mice exhibited diminished trabecular bone quality parameters at 7-months which remained significantly lower than the wild-type mice at 18 months-of-age. ABCC6^-/-^ vertebrae showed increased TRAP staining along with decreased TNAP staining, suggesting an enhanced bone resorption as well as decreased bone formation. Surprisingly, however, loss of ABCC6 resulted only in a mild, aging disc phenotype without evidence of dystrophic mineralization. Finally, we tested the utility of oral K3Citrate to treat the vertebral phenotype since it is shown to regulate hydroxyapatite mechanical behavior. The treatment resulted in inhibition of osteoclastic response and an early improvement in mechanical properties of the bone underscoring the promise of potassium citrate as a therapeutic agent. Our data suggest that although ectopic mineralization is tightly regulated in the disc, loss of ABCC6 compromises vertebral bone quality and dysregulates osteoblast-osteoclast coupling.

**Author Summary:** Inherited mutations in the ABCC6 transporter gene results in mineralization, often in the form of hydroxyapatite, of connective tissues throughout the body, predominantly affecting the skin, eyes, and blood vessels. Functional loss of ABCC6 causes reduced levels of the potent mineralization inhibitor pyrophosphate (PPi) in blood resulting in these pathologies. Pathological mineralization is also a prominent feature of intervertebral disc degeneration, but the role of ABCC6 and systemic PPi levels and its correlation to disc mineralization and vertebral bone health has remained unexplored. In this study, we show for the first time that loss of ABCC6 in mice results in significant decline in vertebral bone quality and mild age-related disc degeneration without increased incidence of abnormal mineralization. Importantly, treatment of ABCC6 deficient mice with K3Citrate resulted in restoration of early cellular changes which drive bone loss and mechanical function of the vertebrae. In summary, our data reveal that ABCC6 is dispensable for mineralization prevention in the intervertebral disc. Unexpectedly, we found that vertebral bone quality and bone cell activities are linked to ABCC6 function.

## Introduction

The ABCC6 protein is a mediator of cellular ATP release [1, 2] into the blood from hepatocytes where it is primarily expressed. ABCC6 inhibits ectopic mineralization, wherein extracellular ATP is converted to AMP and the key mineralization inhibitor PPi via ectonucleotide pyrophosphate/phosphodiesterase 1 (ENPP1). Absence of ABCC6 causes pseudoxanthoma elasticum (PXE), an autosomal recessive metabolic disorder characterized by ectopic mineralization in elastin-rich tissues such as the eyes, blood vessel walls, and the skin [3]. To date, over 300 mutations in the ABCC6 gene have been identified, the majority of which are single nucleotide missense mutations that result in protein function loss [4]. Consequently, global ABCC6 knockout mouse is commonly used to study PXE phenotypes [5, 6]. Noteworthy, while ABCC6 is highly expressed in the liver and kidneys, it is minimally expressed, or absent, in elastin-rich tissues affected by PXE [7]. The PPi produced in liver is distributed to peripheral organs through blood, with ABCC6 contributing about 60-70% of plasma PPi [1, 2] underscoring the metabolic nature of PXE.

Mineral homeostasis is under tight regulation to maintain appropriate development of various tissues including cartilage, bone, and other connective tissues. When mineralization is dysregulated, however, it can result in ectopic mineralization, the aberrant deposition of calcium-phosphate hydroxyapatite crystals in soft connective tissues [8]. Ectopic mineralization affects the connective tissues of the spinal motion segment including the intervertebral disc, facet joint cartilage and ligaments [9]. While disc degeneration is multifactorial, one of the disease sub- phenotypes is ectopic calcification of proteoglycan rich nucleus pulposus (NP) and cartilagenous endplates (CEP) [10, 11], where the latter has been proposed to cause changes to the blood supply and block diffusion of nutrients into the disc [12, 13]. In a recent study using high-resolution imaging digital-contact radiography a strong correlation between aging, disc degeneration status and pathological calcification of intervertebral disc and between disc and facet joint degeneration and facet cartilage calcification has been shown [14]. Noteworthy, 100% disc and 36.5% facet cartilage specimens showed calcifications indicating that pathological calcification is a prominent feature of the degenerative process [14]. Several mouse models such as spontaneous Enpp1asj-2J mutant mice [15], a model of generalized arterial calcification of infancy (GACI) and aging inbred LG/J mice [16] have shown high incidence of disc calcification linked to degeneration that affects the fibrocartilaginous annulus fibrosus (AF) and NP compartments [16]. Other ectopic mineralization disorders such as diffuse idiopathic skeletal hyperostosis (DISH), caused by a lack of equilibrative nucleoside transporter 1 (ENT1), is characterized by increased ectopic mineralization of AF tissue and bone mineral density which progresses with age [17, 18]. These ENT1 null mice show reduced expression of anti-mineralization genes *MgP*, *Enpp1*, and *Spp1* by 6 months of age and enhanced ectopic spine calcification from cervical to caudal regions in an age-dependent manner [19].

Surprisingly, reports of PXE-related calcifications in the musculoskeletal system have only until recently been investigated in zebrafish, where the knockout of the ABCC6 ortholog resulted in hypermineralization of the axial skeleton [20]. A very recent study showed that a small subset, 16.7%, of adult GACI patients with mutations in ABCC6 suffered from hypophosphatemic rickets [21], implying that bone is an affected tissue in relation to ABCC6 mutations. It is also important to note that the intervertebral disc contains an extensive elastin fiber network [22], which increases with aging and degeneration [23]. Consequently, discs could be susceptible to mineralization in the context of PXE spectrum of disorders. Given that PPi is a major inhibitor of connective tissue mineralization, it is unclear whether ABCC6-derived systemic PPi is important for preventing disc calcification.

Here, for the first time we provide insights into the role of ABCC6 in vertebrae and the intervertebral disc in the context of aging using ABCC6 knockout mice. Our results show that loss of ABCC6 significantly affects the vertebral bone morphology and mass. Moreover, we show that K3Citrate could have a potential in treating the observed osteopenic phenotype due to its known propensity to accumulate in bones where it plays an important function in regulating hydroxyapatite mechanical behavior [24]. Interestingly, we observed that ABCC6 loss does not predispose discs to pathological mineralization. These results highlight the significance of local vs. systemic regulation of PPi metabolism in the musculoskeletal system.

## Results

### The ABCC6^-/-^ mice show vertebral osteopenia, alterations in disc height and reduced osteoblast and osteoclast activity

Vertebral bone properties have not been previously shown to be affected in PXE patients or animal models. MicroCT analysis of ABCC6^-/-^ vertebrae, however, showed robust and age-dependent decline in trabecular bone attributes in lumbar (Fig. 1) and caudal (S1 Fig.) spine along with alterations in the disc height (Fig. 1a-h’). Trabecular parameters that were reduced in ABCC6^-/-^ mice included bone volume fraction (BV/TV) (Fig. 1i), trabecular thickness (Tb.Th.) (Fig. 1j), and trabecular number (Tb.N.) (Fig. 1k), with a concomitant increase in trabecular spacing (Tb.Sp.) (Fig. 1l) at all timepoints except at 23-weeks. Structural model index (SMI), a parameter that determines the rod-like structure of trabeculae and is correlated to bone strength and fracture risk, was significantly higher at all timepoints except 23-weeks (Fig. 1m), suggesting that ABCC6^-/-^ trabeculae have decreased bone strength and higher fracture risk. This observation was also supported by an initial higher bone mineral density (BMD) at 23-weeks, but then a progressive reduction in BMD from 7-8-months onward (Fig. 1n). Furthermore, these changes in lumbar trabecular bone in ABCC6^-/-^ mice were confirmed in the caudal spine (S1 Fig. i-n). MicroCT analysis showed less of an impact of absence of ABCC6 on lumbar cortical bone. Only at 12-months was there an increase in bone area (B.Ar.) (Fig. 1o), mean polar moment of inertia (MMI) (Fig. 1p), and tissue mineral density (TMD) (Fig. 1r). Interestingly, cross-sectional thickness (Cs.Th.) was only significantly reduced at 7-8 months and 16-18 months, whereas it trended higher at 12-months (Fig. 1q). Cortical bone in caudal vertebrae of ABCC6^-/-^ was relatively unaffected with a slightly decreased MMI at 12-months and decreased TMD at 23- weeks (S1 Fig. o-r). When assessing the relationship between the disc compartment and vertebrae, ABCC6^-/-^ mice had increased vertebral length and alterations in disc height and disc height index (DHI) (Fig. 1 s-u), parameters correlated with disc degeneration. Caudal vertebral length, disc height and DHI, however, only showed significant differences at 7-8 months (S1 Fig. s-u). Together, these results clearly showed that loss of ABCC6 altered bone mass and morphology and resulted in an age-progressive osteopenic phenotype.

**Fig 1.**
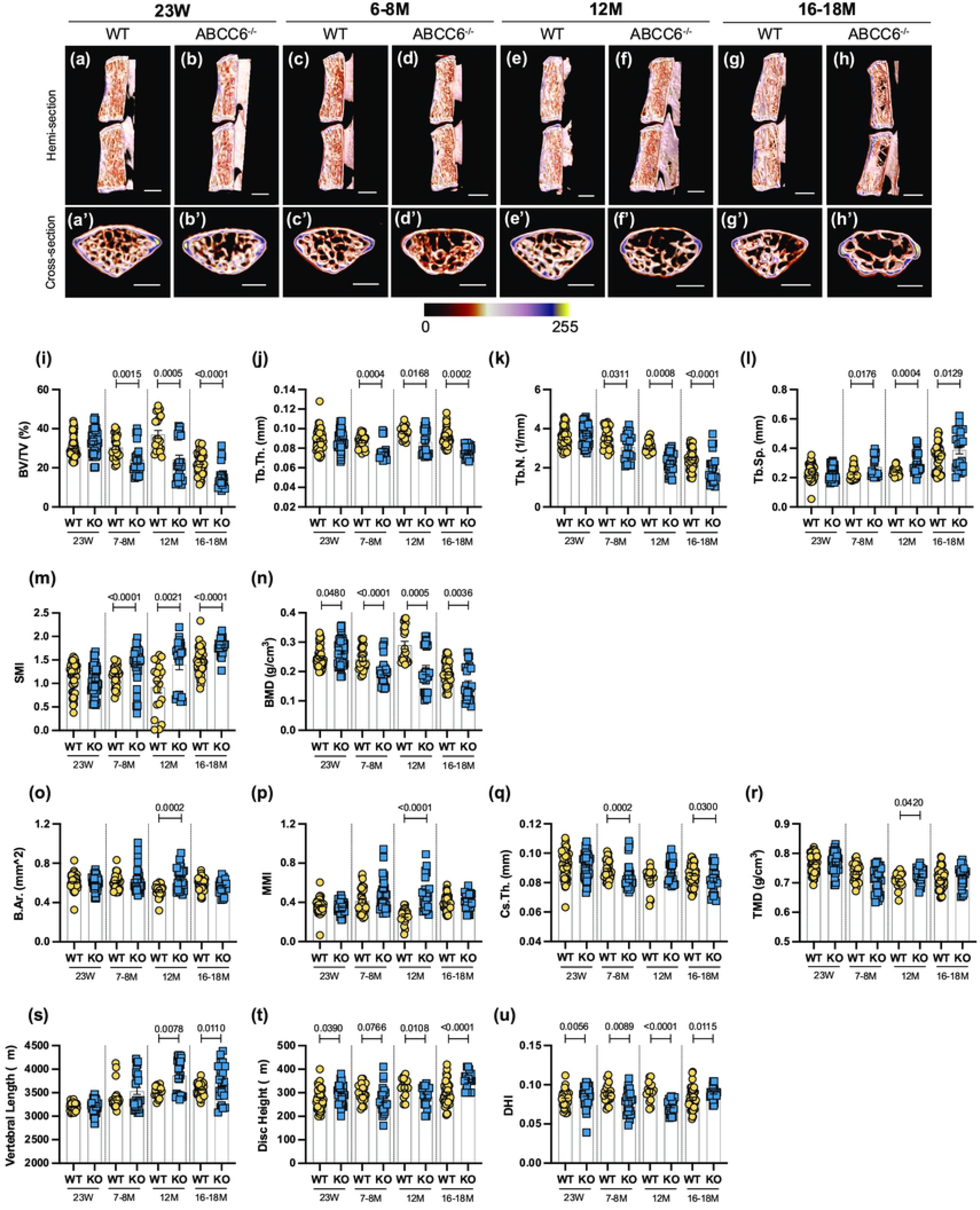
ABCC6^-/-^ mice show vertebral osteopenia, altered disc height and vertebral length. Representative microCT reconstructions of *(a-h)* hemi- and *(a’-h’)* cross-sections show consistent trabecular thinning in lumbar vertebrae of ABCC6^-/-^ mice at all ages. Quantitative microCT analysis of trabecular bone parameters *(i-n)* BV/TV, Tb.Th., Tb.N., Tb.Sp., SMI, BMD, and cortical bone parameters *(o-r)* B.Ar., MMi, Cs.Th., TMD. *(s)* Vertebral length, *(t)* disc height, and *(u)* DHI are shown for lumbar motion segments. Quantitative analyses are shown as mean ± SD (n = 3 lumbar discs and n = 4 vertebrae/mouse, n ≥ 5 mice/genotype). Significance was determined using unpaired t-test or Mann Whitney as appropriate. *(a-h)* Scale bar = 1 mm. *(a’-h’)* Scale bar = 500 μm. BV/TV= bone volume/tissue volume. Tb.Th.= trabecular thickness. Tb.N.= trabecular number. Tb.Sp.= trabecular spacing. SMI = structural model index. BMD = bone mineral density. B.Ar.= bone area. MMI= mean polar moment of inertia. Cs.Th.= cross-sectional thickness. DHI= disc height index.

Interestingly, ABCC6^-/-^ vertebrae showed a significant increase in TRAP staining, associated with the presence of osteoclasts (Fig. 2 a-c), and a concomitant decrease in TNAP levels (Fig. 2 d-f) at 7-8 months of age. This initial dysregulation in osteoblastic activity is in line with the observed morphometric deficits suggesting that upregulated osteoclastic activity is driving this progressive trabecular bone loss. These changes, however, were not maintained at 16-18 months with TRAP expression showing only a trend of increased activity over age-matched wild-type mice, implying that over time the loss of bone is maintained due to decoupling of bone formation and resorption.

**Fig 2.**
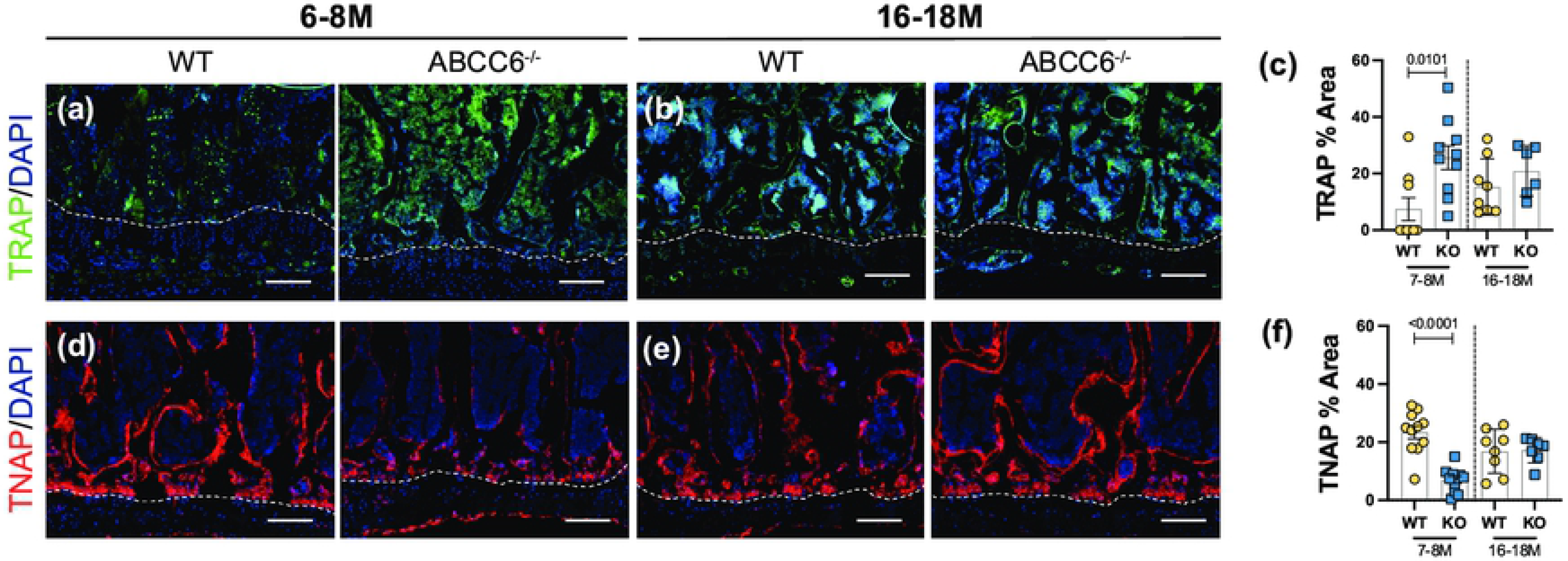
Osteoblastic activity is dysregulated in lumbar vertebrae of ABCC6^-/-^ mice. Immunohistological (IHC) staining showed increased *(a-b)* TRAP staining and decreased *(d-e)* TNAP staining in 7–8-month ABCC6^-/-^ lumbar vertebrae. No difference between genotypes was observed in TRAP and TNAP activity in 16–18-month mice. Scale bar = 100 μm. Quantitative analyses are shown as mean ±SD. (n = 2 vertebrae/mice, n ≥ 5 mice/genotype/stain). Significance was determined using unpaired t-test or Mann Whitney test as appropriate.

### Loss of ABCC6 causes small changes in cellular phenotype but does not promote age-dependent disc degeneration or mineralization

Although there are numerous reports on ectopic mineralization of elastin-rich tissues resulting from ABCC6 loss-of-function, there have been none that investigate the significance of this loss on the intervertebral disc. Histological analysis did not show conspicuous changes in lumbar NP and AF cell morphology (Fig. 3a-b’). A Modified Thompson grading scheme of NP and AF tissue revealed age-related disc degeneration in both wild-type and ABCC6^-/-^ mice, particularly in the AF, a higher proportion of discs scored a grade 3 in the AF at 16-18 months compared to WT discs (Fig. 3c). On the other hand, ABCC6^-/-^ at 16-18 months showed fewer discs with grade 2 NP (Fig. 3d).

**Fig 3.**
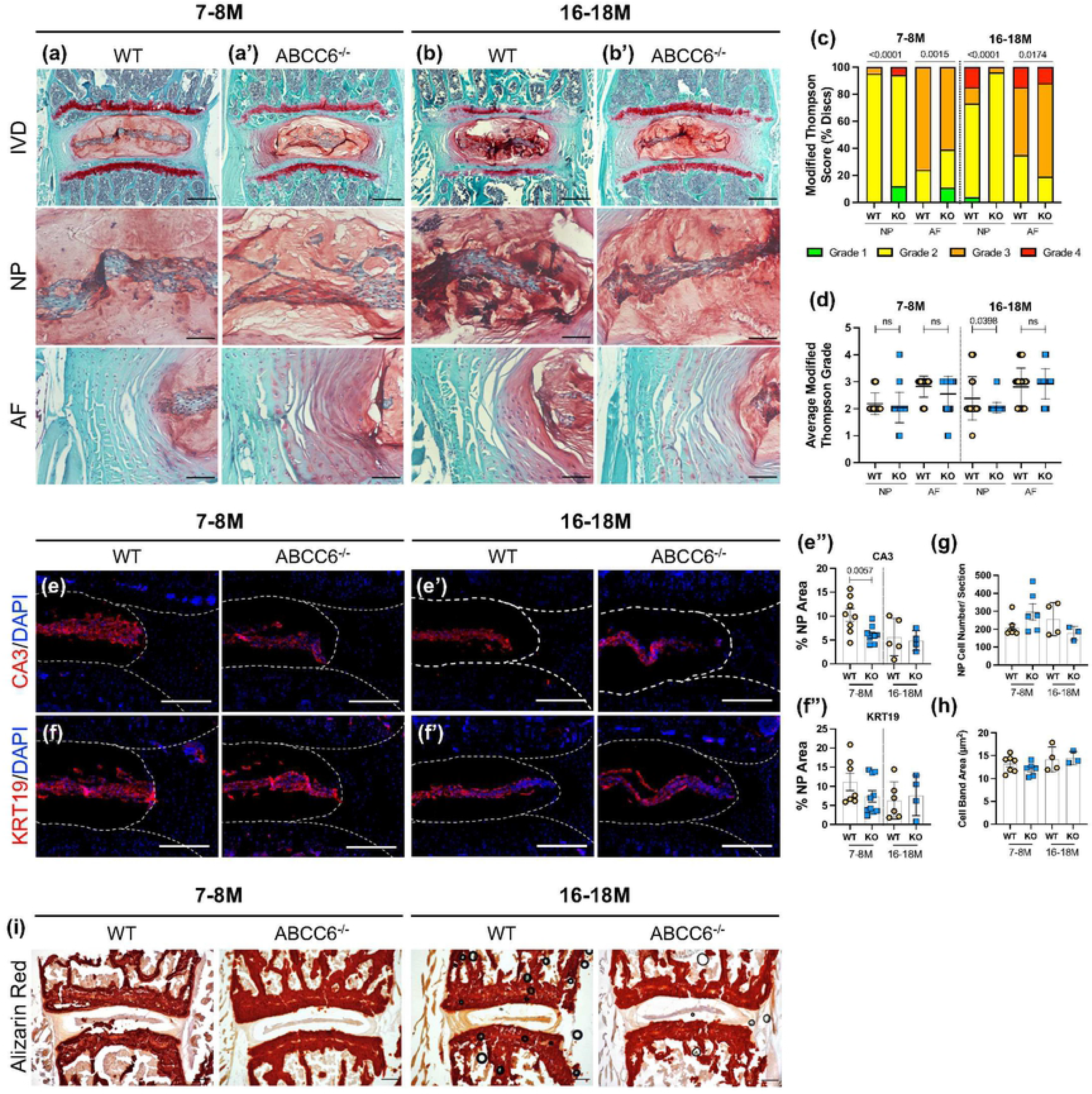
ABCC6 loss shows mild degenerative changes but does not promote disc mineralization. (a-b’) Safranin O/Fast Green staining of (a-a’) 7-8 month and (b-b’) 16–18- month-old lumbar discs showed tissue morphology and proteoglycan content consistent with age-related disc degeneration (row 1, scale bar = 200μm and rows 2-3, scale bar = 50μm). *(c, d)* Histological grading analysis using the modified Thompson scale showed changes in distribution of grades of degeneration of NP and AF but comparable average grades of degeneration. *(e-f”’)* IHC staining of NP phenotypic markers of 7-8- and 16-18-month-old lumbar discs showed a slight decrease in *(e-e”)* CA3 expression but no difference in *(f-f”)* KRT19 expression. *(g)* Average NP cell count and *(h)* average NP cell band area showed no difference between genotypes. *(i)* Representative Alizarin Red staining of 7-8- and 16-18-month-old lumbar discs showed no changes in free calcium staining within the NP or AF. Scale bar = 200 μm. Dotted lines denote NP and AF tissue compartments. Quantitative analyses are shown as mean ±SD. Significance of grading distribution was determined using a χ^2^ test. Significance of average grade data and percent area were determined using unpaired t-test or Mann Whitney test as appropriate.

To investigate the phenotype of NP cells of ABCC6^-/-^ mice, levels of phenotypic markers carbonic anhydrase 3 (CA3) and keratin-19 (KRT19) were measured. There was a reduction in CA3 abundance at 7-8 months (Fig. 3 e-e”) with a similar trend in KRT19 levels (Fig. 3 f-f”). However, this early reduction in marker abundance was not sustained as discs aged, suggesting that loss of ABCC6 does not exacerbate age-related decline in cell number and their phenotype (Fig. 3 g-h). In addition, Alizarin red staining was performed to assess calcium levels. Analysis revealed presence of AF calcification in one 16-18M ABCC6^-/-^ mouse (S2Fig. a), however, there was no overall apparent elevation in calcium levels or presence of calcified nodules in ABCC6^-/-^ discs (Fig. 3 i), suggesting that discs are not susceptible to pathological mineralization following the loss of ABCC6.

### Loss of ABCC6 causes alteration in collagen fiber thickness and composition in AF

We performed Picrosirius red staining followed by polarized imaging of WT and KO lumbar discs to investigate changes in AF collagen fiber thickness (Fig. 4a-b’). The analysis of polarized images showed that lumbar discs of ABCC6^-/-^ mice showed an increased proportion of thin collagen fibers and a lower proportion of thick fibers at 7-8 months of age compared to WT discs (Fig. 4 a’, c). Interestingly, these percentages were reversed at 16-18 months with more thin fibers compared to thick fibers, suggesting that loss of ABCC6 causes a disruption in collagen turnover and homeostasis. A similar trend in collagen fiber thickness was observed in caudal discs (S2 Fig. e-g). To gain further insights into compositional changes in collagen subtypes, levels of collagen-I (COLI), collagen-II (COLII), and collagen-X (COLX) were measured. Interestingly, abundance of COLI (Fig. 4 d-d”) and COLII (Fig. 4 e-e”) in the AF compartment were comparable between genotypes, suggesting that the changes in proportion of fiber types were not due to COLI and COLII and that other fibrillar collagens could be involved. Notably, levels of COLX, a marker of hypertrophic chondrocytes in the growth plate, were significantly elevated in both the NP and AF of 16-18 month and higher in the NP of 7–8-month ABCC6^-/-^ discs (Fig. 4 f- f”’). Overall, these results show that ABCC6 loss results in changes in collagen configuration that may reflect mild disc degeneration.

**Fig 4.**
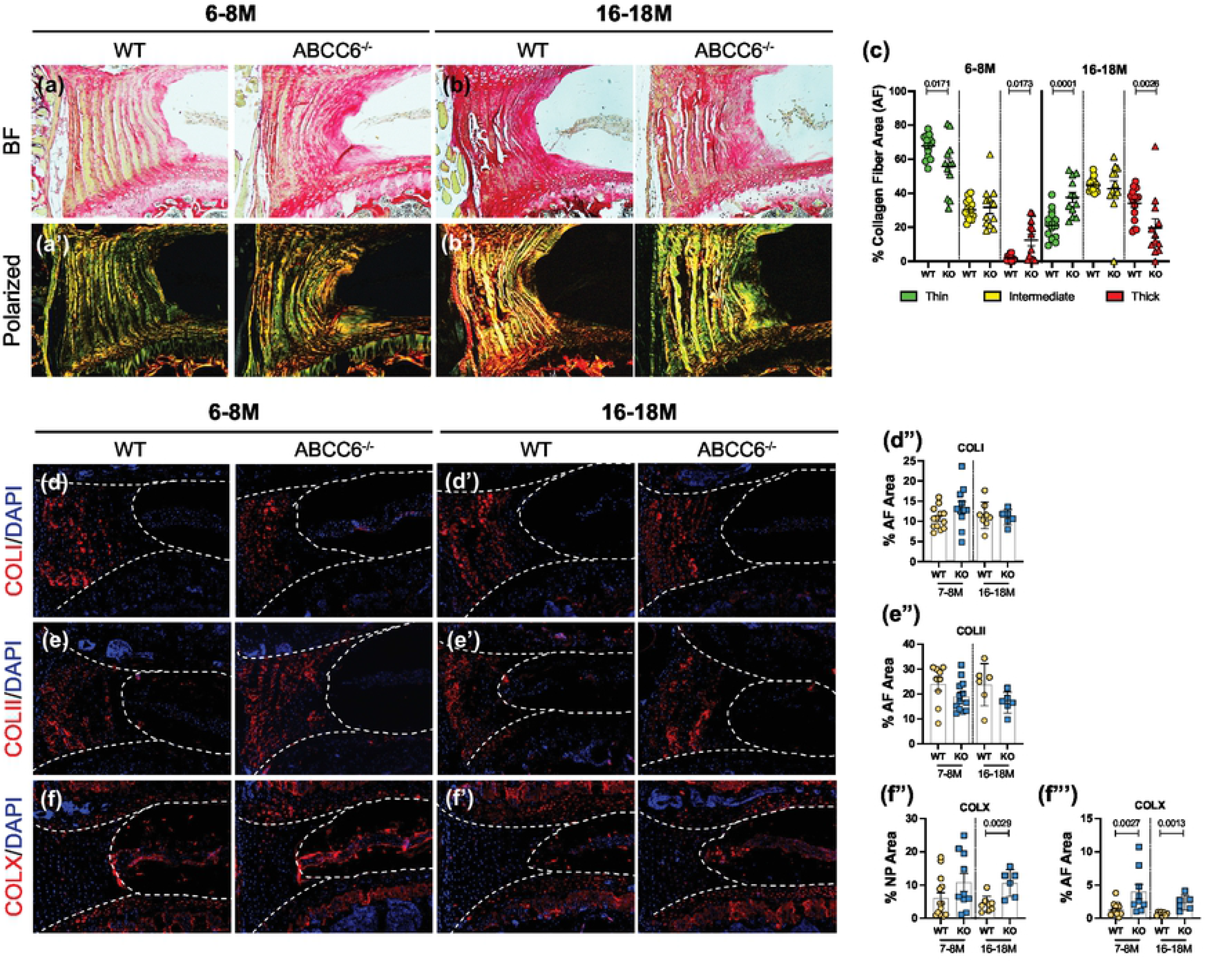
ABCC6^-/-^ mice show altered AF collagen fiber composition. *(a-b’)* Picrosirius Red staining showed altered collagen fiber thickness in the AF of ABCC6^-/-^ lumbar discs under bright field (BF) and polarized light. *(c)* Quantification of percent collagen fiber area showed a significant difference in distribution of fiber maturity. *(d-f”’)* IHC staining showed no differences in *(d, d”)* COLI and *(e, e”)* COLII abundance but increased *(f, f”’)* COLX expression at both timepoints of ABCC6^-/-^ AF tissue. Dotted lines denote NP and AF tissue compartments. Scale bar = 200μM. (n = 1-2 discs/animal; n ≥ 3 mice/genotype, 5-14 discs/genotype/stain) Quantitative analyses are shown as mean ±SD. Significance was determined using unpaired t-test or Mann Whitney test as appropriate.

### ABCC6^-/-^ intervertebral discs show minor compositional changes in non-collagenous matrix components

The impact of ABCC6 deletion on composition of major, non-collagenous ECM components of disc was assessed. Aggrecan (ACAN), a high molecular weight, chondroitin sulfate (CS) and keratan sulfate substituted proteoglycan, was highly expressed in the NP compartment and showed increased abundance in 16-18-month-old ABCC6^-/-^ mice; comparable levels in NP were noted between genotypes at 7-8 months (Fig. 5 a-a”’). There were no differences in aggrecan levels in the AF compartment between KO and WT mice at both time points. While no changes were seen in the NP compartment, levels of CS were reduced in the AF of 16–18-month ABCC6^-/-^ discs with a trend of decreasing levels at 7-8 months (Fig. 5 b- b”’). Staining for ARGxx, an ACAN neoepitope generated by ADAMTS-dependent degradation, showed a small trend of increase in the AF compartment of ABCC6^-/-^ mice at 7-8 month, whereas a significant decrease was noted at 16-18 months. (Fig. 5 c-c”’). Furthermore, cartilage- oligomeric matrix protein (COMP), another important non-collagenous matrix component, showed comparable abundance across ages and genotypes (Fig. 5 d-d”’). Taken together, these results suggest alteration in turnover and CS-substitution of ACAN in the AF with concomitant compensatory increase in ACAN levels in the NP of ABCC6^-/-^ mice.

**Fig 5.**
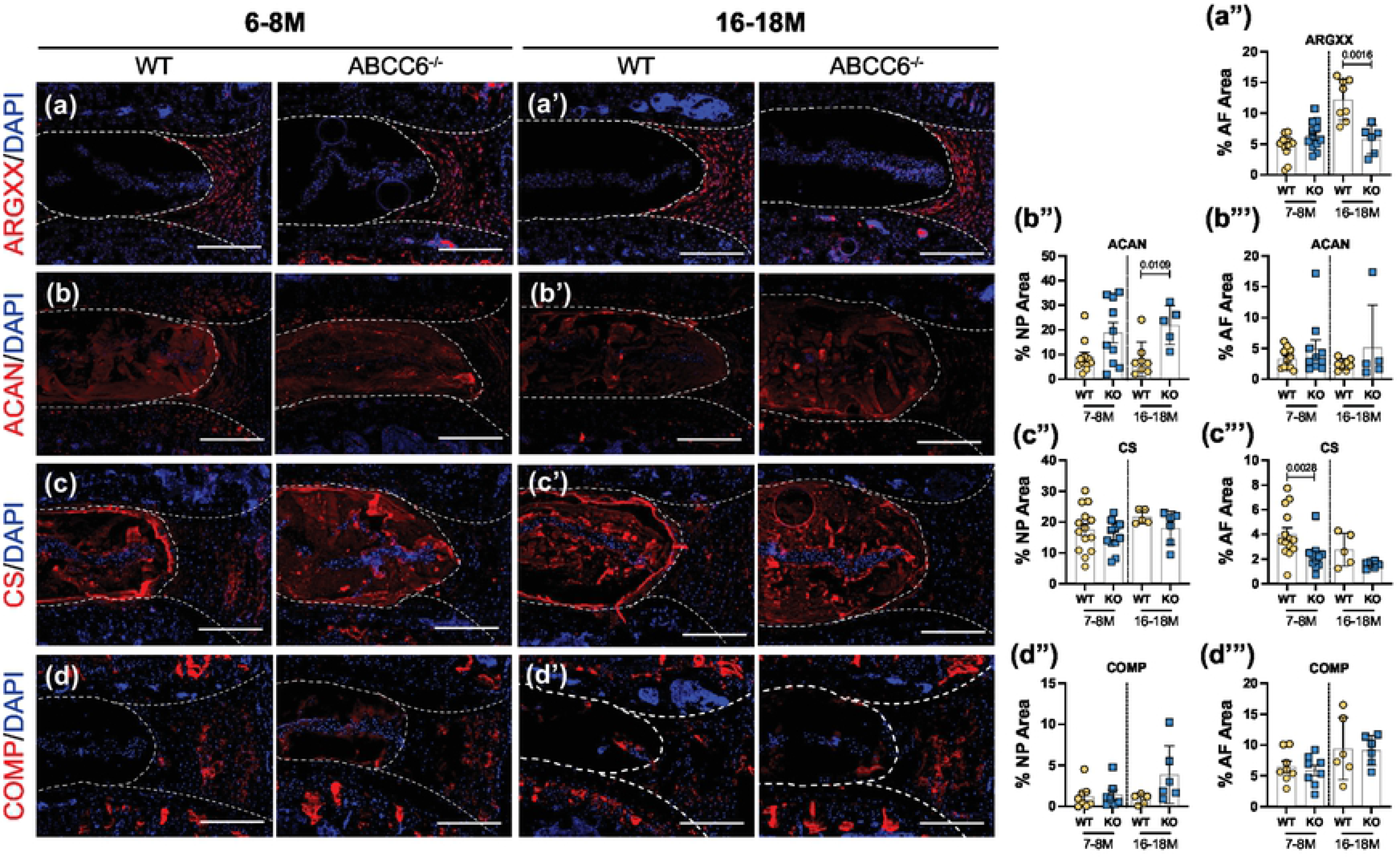
ABCC6^-/-^ mice exhibit changes in non-collagenous disc matrix composition. *(a-d’)* IHC staining showed reduced expression of *(a, a”)* ARGxx and *(c, c”’)* CS in the AF, increased expression of *(b, b”’)* ACAN in the NP, and no changes in *(d, d”’)* COMP expression in 16–18- month lumbar discs. Scale bar = 200μM. (n = 1-2 discs/animal; n ≥ 3 mice/genotype, 5-14 discs/genotype/stain) Quantitative analyses are shown as mean ±SD. Significance was determined using unpaired t-test or Mann Whitney test as appropriate.

### ABCC6^-/-^ mice reveal transcriptomic changes related to metabolic processes and sensory perception

Since AF tissue has extensive elastin network and undergoes ectopic mineralization in humans and mouse models of disc degeneration, we performed microarray analysis of AF tissue from 7-month-old mice with an aim to obtain molecular and mechanistic insights into early transcriptomic changes resulting from ABCC6 loss. This age also corresponded with emergence of an osteopenic phenotype in vertebrae. Three-dimensional Principal Component Analysis (PCA) showed distinct clustering of WT samples and consistent clustering among three of the four KO samples (Fig. 6 a). At p ≤ 0.01 and 1.75-fold-change threshold, there were a total of 144 differentially expressed genes (DEG’s) in AF (Fig. 6 b), where hierarchical clustering heatmap and volcano plot show the directionality and proportions of DEG’s (Fig. 6 c, d).

**Fig 6.**
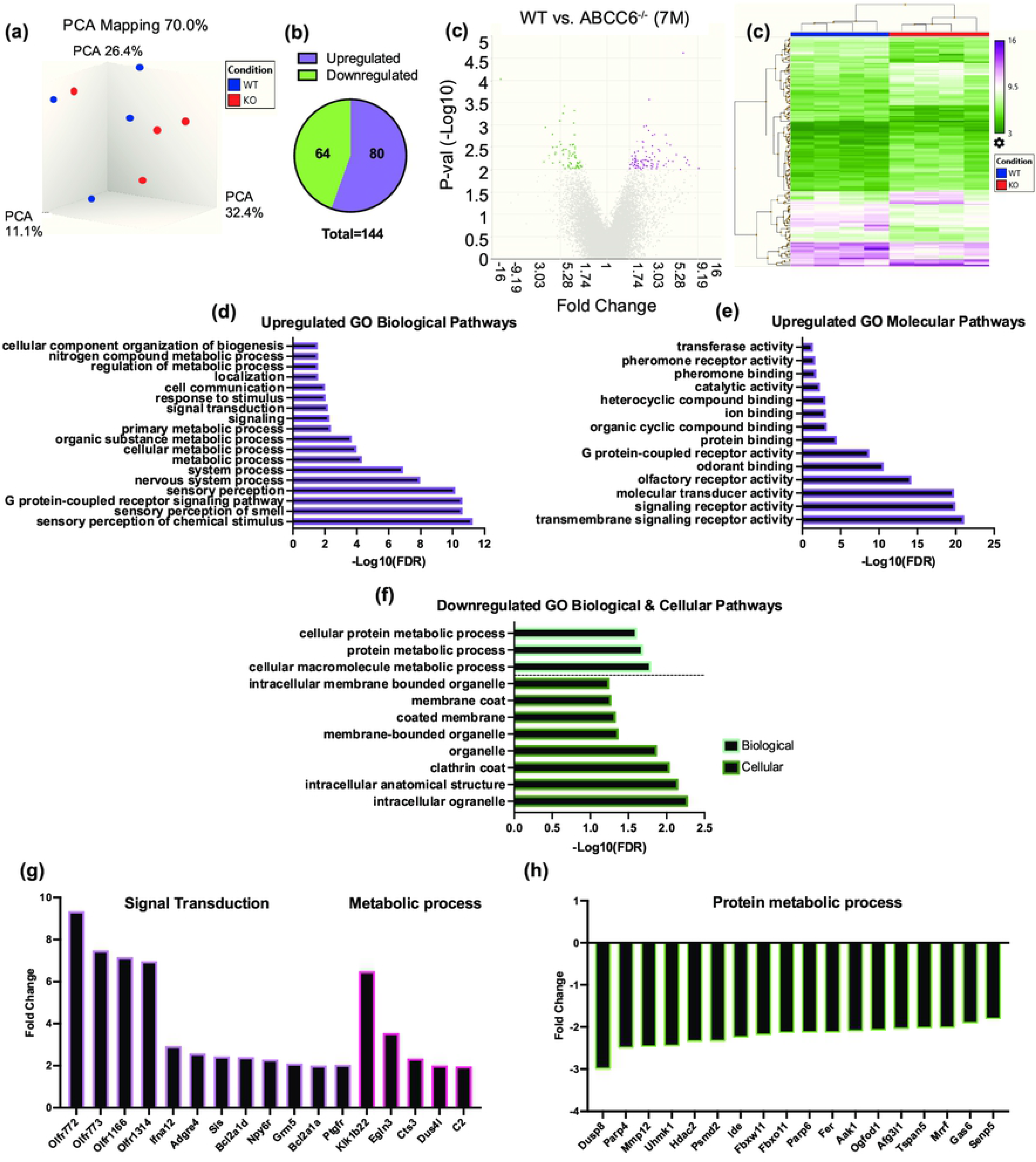
Microarray analysis shows minimal differential expression of genes in ABCC6^-/-^ mice in AF tissue. *(a)* Principal Component Analysis showed clustering of samples. *(b)* Venn diagram showed a total of 144 differentially expressed genes (DEGs) and *(c)* volcano plot (p < 0.01, ≥ 1.75-fold change). *(d)* Hierarchical clustering showed the directionality of significant DEGs. Upregulated GO *(e)* Biological Processes (BP) and *(f)* Molecular Function pathways. *(g)* Downregulated GO BP and GO Cellular Component pathways. Representative *(i)* upregulated and *(j)* downregulated DEGs from select GO processes.

Enrichment analysis was performed using PANTHER to assess GO biological, molecular, and cellular pathways related to these DEGs. From this analysis, among the most enriched upregulated GO Biological and Cellular processes included sensory perception, cellular metabolic process, signal transduction, G protein-coupled receptor activity, and ion binding (Fig. 6 e, f). Among the most enriched downregulated GO processes included cellular macromolecule molecular process, protein metabolic process and cellular processes related to clathrin coat and membrane-bound organelle (Fig. 6 g). The select genes classified under these enriched pathways included upregulation of signal transduction genes *Ifna12*, *Npy6r*, *Bcl2a1d*, and *Ptgfr* and metabolic process genes *Klk1b22* and *Egln3*. Whereas downregulated DEGs included *Mmp12*, *Hdac2*, and *Tspan5* (Fig. 6 i, j).

### K3Citrate treatment restores early cellular changes and mechanical performance in ABCC6^-/-^ vertebrae

We investigated whether oral supplementation K3Citrate can reverse the early cellular and mechanical changes which may occur prior to the onset of bone loss. We analyzed vertebrae from 23-weeks old ABCC6^-/-^ mice that were treated with vehicle or K3Citrate at 2 mM and 40 mM for 20 weeks starting at 3-weeks of age. While 23-week-old ABCC6^-/-^ mice do not evidence bone loss, they showed increased TRAP abundance compared to WT animals. This increased TRAP levels in KO vertebrae was restored to WT levels in cohorts treated with 2mM K3Citrate and showed a further reduction at 40mM K3Citrate (Fig. 7 a-e). Unlike altered TRAP labeling, at this age, vertebral TNAP activity in ABCC6^-/-^ was comparable to WT. However, vertebral TNAP abundance was significantly increased in ABCC6^-/-^ mice treated with 40mM K3Citrate (Fig. 7 f-j) suggesting stimulation of osteoblastic function. These results suggest that bone loss from ABCC6 deletion was initiated by elevated osteoclast activity which was subsequently followed by osteoblastic dysfunction.

**Fig 7.**
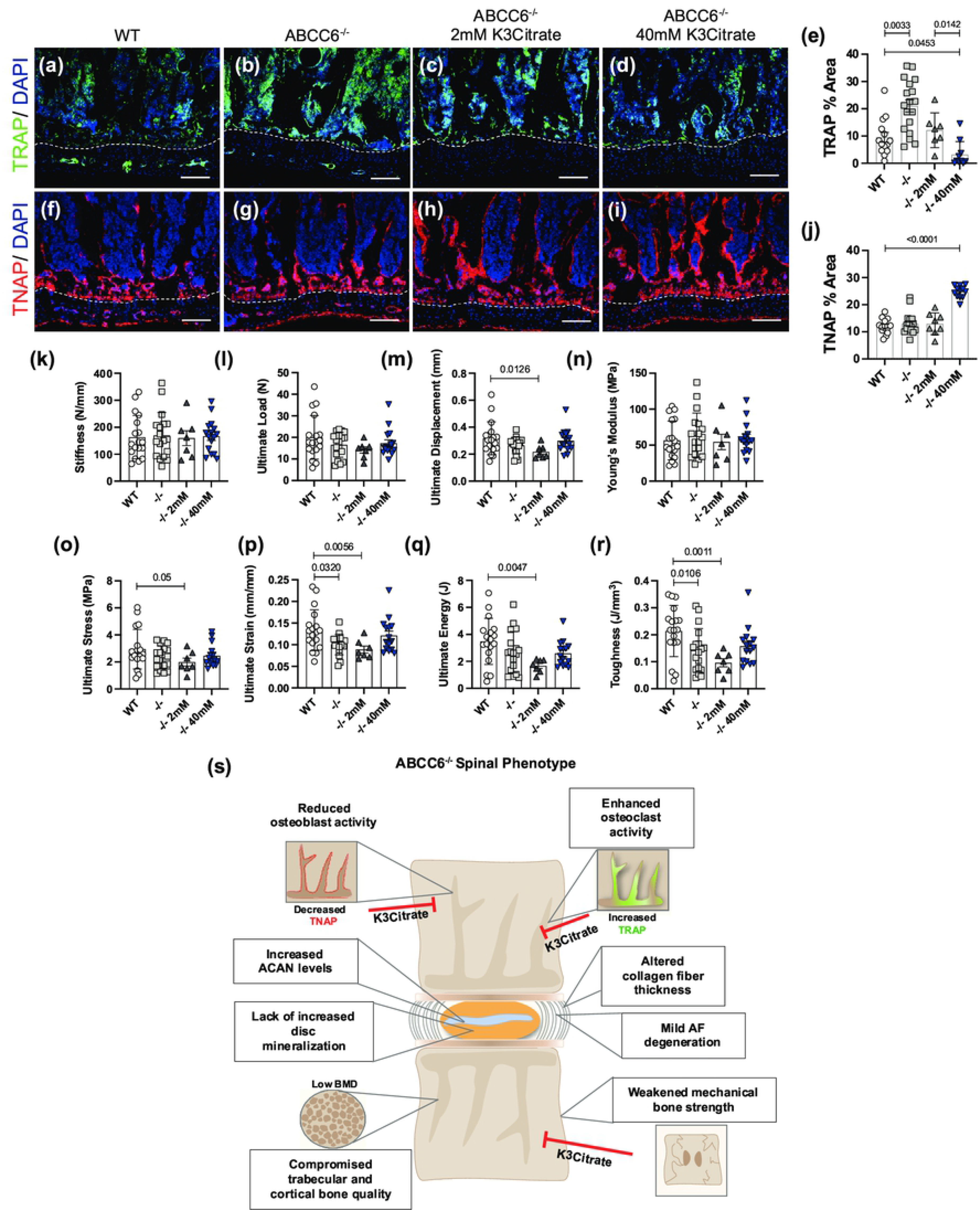
Potassium Citrate (K3Citrate) treatment blocks osteoclast activation, increases osteoblastic activity, and restores mechanical strength of 23-week-old ABCC6^-/-^ lumbar vertebrae. IHC staining showed decreased *(a-e)* TRAP staining and increased *(f-j)* TNAP staining in ABCC6^-/-^ lumbar vertebrae after 40mM K3Citrate treatment. *(k-r)* Biomechanical analysis revealed a restoration of *(p)* ultimate strain and *(r)* toughness in ABCC6^-/-^ lumbar vertebrae after 40mM K3Citrate treatment. *(s)* Schematic representing the role of ABCC6 loss on the spinal motion segment and the effects of K3Citrate treatment. Scale bar = 100μm. Quantitative analyses are shown as mean ±SD. (n = 2 vertebrae/mice, n ≥ 4 mice/group). Significance was determined using ANOVA or uncorrected Kruskal-Wallis test as appropriate.

Biomechanical analysis of lumbar vertebrae from 23-week-old mice revealed that vertebrae of ABCC6^-/-^ evidence lower ultimate strain (Fig. 7 p) and toughness (Fig. 7 r), suggesting that they experience hampered deformation capability and reduced energy absorption before fracture. Importantly, this reduced bone strength in ABCC6^-/-^ vertebrae was restored when mice were supplemented with 40mM K3Citrate. There were no differences in stiffness (Fig. 7 k) and ultimate load (Fig. 7 l) between WT and ABCC6^-/-^ mice or the treatment groups. However, although not statistically significant, similar trends were observed for ultimate displacement (Fig. 7 m), ultimate stress (Fig. 7 o), and ultimate energy (Fig. 7 q), suggesting that reduced biomechanical strength of vertebrae in ABCC6^-/-^ mice can be rescued with K3Citrate treatment.

Overall, these data suggest that early treatment of K3Citrate improved the cellular and mechanical changes that preceded morphological and structural changes to the bone. Therefore, long term therapeutic treatment of K3Citrate has the potential benefit of abating trabecular and cortical bone loss by rescuing the mechanical deficit and reverting the cellular changes observed in the vertebrae. Figure 7s summarizes the spinal phenotype of ABCC6^-/-^ mice and highlights the potential of K3Citrate treatment to treat the vertebral defects.

## Discussion

PXE is an extensively studied mineralization disorder and use of mouse models recapitulating PXE have provided extensive knowledge about ABCC6 function in maintaining the health of elastin-rich tissues that are prone to ectopic mineralization. While dystrophic calcification is one of the known phenotypes of intervertebral disc degeneration, it is unknown whether PXE affects mineralization pathways in the disc and although ABCC6 is not expressed in the spine, its systemic effects on the spine have not been studied in detail. In ABCC6^-/-^ mice we found no increase in calcification of the disc, but unexpectedly, that these animals had reduced vertebral bone quality.

There are a limited number of previous studies of patients and mouse model of PXE that have investigated long bone phenotype. A recent clinical study reported that PXE patients have a higher OA score in knee and articular joints [25], however no effects on long bone mineral density in PXE patients and ABCC6 knockout mice has been noted [26–28]. This study for the first time observed an osteopenic phenotype in the axial skeleton through a significant reduction of vertebral bone morphology parameters, including reduced trabecular bone volume, thickness, and number as well as reduced bone mineral density. These findings are in line with the early onset osteoporosis and spinal fractures seen in adult patients with heterozygous loss-of-function mutations in ENPP1 and osteoporosis in ENPP1^-/-^ mice, a model for GACI [29]. Likewise, reduced BMD in lumbar spine has been noted in ENT1^-/-^ mice, a model for diffuse idiopathic skeletal hyperostosis (DISH)^2^ [29, 30]. These findings suggest that the axial skeleton is sensitive to the impairment of PPi levels. Furthermore, changes in bone turnover markers, such as increased TRAP and reduced TNAP abundance, have been associated with disc degeneration and osteoporosis and are considered valuable tools for understanding the cellular basis of bone loss [31, 32]. The elevated TRAP abundance observed before the structural deficit in vertebral bone was evident, suggests that enhanced osteoclastogenesis in ABCC6^-/-^ bone is the driving mechanism for such bone loss and that osteoblastic dysregulation evident by reduced TNAP staining, follows osteoclastic activation.

Histological analyses at various ages provided evidence of mild but age-progressive disc degeneration in the AF compartment of ABCC6^-/-^ mice. Aging promotes degeneration in both the NP and AF compartments [16], however only the AF of 16-18-month ABCC6^-/-^ exhibited a higher incidence of degeneration. While decreased abundance of phenotypic markers CA3 and KRT19 is consistent with aging and degeneration [33, 34], yet the minor, or lack of, progressive changes observed in ABCC6^-/-^ mice beyond 7-8 months suggested overall preservation of NP function. Importantly, the lack of major morphological changes in the NP compartment in ABCC6^-/-^ mice explains absence of severe deterioration in the overall architecture of the discs.

While changes in the ABCC6^-/-^ disc were relatively mild, AF compartment appeared to be more sensitive to absence of ABCC6, possibly because of the systemic reduction in extracellular PPi levels.

Despite the mild phenotypic differences, transcriptomic analysis showed enrichment of GO processes related to altered signaling and metabolic processes. Of these GO processes, the genes Npy6r, Ptgfr, and Egln3 were upregulated. Npy6r expression is correlated with disc degeneration and has shown to increase expression in AF cells when exposed to inflammatory stimuli [35, 36]. While Ptgfr blocks the interaction of prostaglandin F2-alpha (PGF2α) to its receptor, Egln3, or PHD3, modulates NF-ĸB and HIF-1α signaling pathways and is controlled by both HIF-1α and proinflammatory cytokines [37–39]. Likewise, expression of MMP12, a promoter of ECM degradation and disc fibrosis [40, 41], Hdac2 and Tspan5, modulators of RANKL-induced osteoclastogenesis [42, 43] were all downregulated. Overall, microarray analysis suggested elevation in inflammatory response genes in the AF of ABCC6^-/-^ discs.

Studies of human disc specimens and mouse models have shown a strong correlation among disc calcification, aging and degeneration status [14,16,17,44], suggesting that dystrophic calcification is one of important sub phenotypes of the degenerative process. ABCC6 null mice experience mineralization in elastin- and/or collagen rich connective tissues of skin, eyes and arteries [6] and ENPP1 null mice show hypermineralization of collagen-rich tendons and ligaments [15], suggesting that the elastin and collagen-rich AF tissue of the disc could be a potential site of ectopic mineralization in ABCC6^-/-^ mice. A notable observation in our study was that there was no drastic increase in incidence of calcification within the NP and AF compartments of aged ABCC6-/- mice. These results suggest that in the avascular environment of the disc, calcium and PPi levels are tightly regulated regardless of the changes in systemic levels of PPi. Relevant to this phenotype, maintenance of extracellular matrix components that may contribute to modulation of mineralization is critical for disc function. For example, COMP, a marker for healthy and young discs is prominently localized in the AF compartment [45] and shown to be an inhibitor of vascular calcification [46], and the proteoglycan aggrecan inhibits apatite growth through Ca^2+^ binding [47]. Consequently, comparable COMP and ACAN levels in the AF accompanied by decrease in ACAN turnover and elevated ACAN levels in NP, may suffice to prevent disc mineralization in ABCC6^-/-^ mice. It is also important to note that COLX, a protein highly expressed by hypertrophic chondrocytes [10, 48] is also a regulator of mineralization in the growth plate through its inhibitory actions and is elevated in ABCC6^-/-^ mice [49].

Considering the importance of balancing the Pi/PPi ratio in regulating connective tissue mineralization, it would be expected that a decrement in systemic levels of PPi will result in ectopic mineralization. Ectopic mineralization, however, might be governed by the local vs. systemic regulation of PPi in a tissue-dependent fashion. The tissues usually affected in PXE patients are more reliant on, and therefore more susceptible to, systemic plasma concentrations of PPi. This point is underscored by a recent study comparing the phenotypes of patients with ENPP1 and ABCC6 deficiencies [21]. The authors reported a significantly lower incidence of joint calcification in ABCC6-deficient patients than patients with ENPP1-deficeit, suggesting that local regulation of PPi and AMP levels may be critical in ectopic mineralization of cartilagenous tissues [21]. The intervertebral disc, like articular cartilage, is avascular and utilizes the microvasculature of the cartilagenous endplates (CEPs) for metabolite diffusion and likely relies upon locally generated PPi. Previous studies have shown an association between aortic calcification and intervertebral disc height loss in humans [50]. Disc degeneration also results from nicotine-induced vascular hypertrophy, suggesting that vascular function is important for disc health [51]. Although we did not investigate the morphology and mineralization status of CEP vasculature, it can be presumed based on the mild disc phenotype observed that endplate vascular calcification is not a pathology in this model. Notably, progressive ankylosis protein (ANKH), another critical transmembrane protein responsible for regulating mineralization is highly expressed in osteoblasts, skeletal muscle as well as disc cells [52–54]. ANK*^ank/ank^* mice exhibit a 30% reduction plasma PPi, a 65% reduction in plasma citrate levels and altered long bone geometry along with progressive joint and disc calcification [55–58]. Furthermore, *in vitro* studies revealed that inhibition of ANKH via TNF increases NP cell mineralization^59^ and ANKH levels are elevated, possibly as a compensatory mechanism in degenerated NP tissues from human patients [54], suggesting it may be the predominant player in controlling disc calcification. Therefore, mechanistic investigations of discs from ANK*^ank/ank^* mice would be important to understand its tissue-specific function and causal effect in disc degeneration and calcification and may provide insights into the key role of locally generated PPi.

Current therapeutic interventions to address ectopic mineralization in disorders such as PXE involve a variety of approaches including PPi supplementation [60], bisphosphonates [61], statins [62], TNAP inhibition [63], and dietary supplement of magnesium and citrate [64, 65] Citrate plays an important role in cellular metabolism and about 90% of the body’s source of mobilizable citrate is stored in the bone [66], where it chelates calcium and stabilizes apatite nanocrystals for bone formation. In addition to export of citrate via ANK, osteoblasts are responsible for the uptake, and storage of citrate, where uptake of plasma citrate relies on sodium-dependent plasma membrane citrate transporter SLC13A5 [67]. Furthermore, reduced bone and plasma citrate levels have been associated with osteoporosis and positively correlate with lumbar BMD [68]. ABCC6^-/-^ mice treated with K3Citrate exhibited a dose-dependent decrease in TRAP levels, where a low dose normalized TRAP levels to that seen in wild-type vertebrae and a higher dose resulted in further reduction. Although TNAP activity was comparable in 23-week-old WT and untreated ABCC6^-/-^ mice, TNAP activity significantly increased after 40mM K3Citrate treatment. Taken together, these data agree with previous findings where cultures of osteoclasts have shown a dose-dependent reduction of osteoclastogenesis after K3Citrate treatment [69] and suggests that citrate plays a role in regulating both osteoclast differentiation and possibly osteoblast maturation/function. Furthermore, compression analysis of ABCC6^-/-^ vertebrae revealed compromised structural and material properties, seen via reduced ultimate strain and toughness showing that the vertebrae are more prone to fracture. After 40mM K3Citrate treatment, however, there was a restoration of these parameters. This suggests that long-term treatment of K3Citrate is safe and would provide positive outcome in rescuing vertebral bone quality by promoting direct incorporation of citrate in bone matrix and/or modulating function of bone cells. Moreover, this treatment modality could benefit not only PXE-related changes in vertebrae, but for other conditions that afflict the spine.

In summary, this study provides novel insights into role of ABCC6 and contribution of systemic PPi in the maintaining health of the intervertebral disc and axial skeleton.

## Materials and Methods

### Mice

All animal experiments were conducted following institutional Animal Care and Use Committee (IACUC) protocols approved by Thomas Jefferson University. 23-week, 7–8-month, 12 months, and 16–18-month-old C57BL6J mice with global deletion of ABCC6, a model for PXE, were analyzed [6]. After weaning, 23-week-old mice treated with potassium citrate (K3Citrate) received either 2mM or 40mM orally through drinking water for 20 weeks.

### Micro-CT analysis

Micro-CT scans (Bruker SkyScan 1275) of fixed spines were conducted under parameters 50kV voltage, 200 μA current, 15μm voxel size resolution. Image reconstruction was performed using NRecon Reconstruction software. Three equidistant points were obtained to measure vertebral length and disc height in DataViewer and DHI was calculated as reported before [70]. CTan software was used for 3D data analysis of trabecular bone and to assess bone volume fraction (BV/TV), trabecular thickness (Tb.Th.), trabecular number (Tb.N.), trabecular spacing (Tb.Sp.), structural model index (SMI), and bone mineral density (BMD). 2D analysis of cortical bone assessed bone area (B.Ar.), cross-sectional thickness (Cs.Th.), mean polar moment of inertia (MMI), and tissue mineral density (TMD).

### Histological analysis

Spines were dissected and fixed in 4% PFA for 48 hr., decalcified in 20% EDTA for 15 days and lumbar (L3-L6) and caudal (Ca1-Ca6) discs were embedded in paraffin. Samples used for calcified sections (L1-L3 & Ca6-Ca12) were placed in 30% sucrose solution for 36 hr. immediately following fixation, then embedded in OCT and snap frozen. 7μm thick mid-coronal sections were prepared and stained with 1% Safranin-O, 0.05% Fast Green, and 1% Hematoxylin and Picrosirius Red for morphological assessment and collagen localization respectively. 10μm thick sections prepared for calcified samples were collected using the Tape Stabilized Tissue Sectioning method and Chitosan adhesion method for slide adhesion [71], and stained with Alizarin Red to detect free calcium and mineralized structures. Slides were imaged using Axio Imager 2 microscope (Carl Zeiss) equipped with Axiocam 105 colour camera with a 5x/0.15 N-Achroplan objective and Zen2^TM^ software (Carl Zeiss), and Eclipse LV100 POL polarizing microscope with a 5x/0.15 objective (Nikon) and NIS Elements Viewer software. Gross histology was analyzed using a Modified Thompson grading scale and the scores of n ≥ 4 blind graders were obtained. Grading was performed for 3 lumbar and 5 caudal discs per mouse [23W (*n=8 WT, 8 KO*), 128 discs total; 7-8M (*n=7 WT, 5 KO),* 96 discs total; 12M (*n=5 WT, 6 KO),* 80 discs total; 16-18M (*n=8 WT, 8 KO),* 128 discs total].

### Immunohistology

Mid-coronal sections were deparaffinized and rehydrated using an ethanol series. Following appropriate antigen retrieval, sections were blocked in the appropriate blocking solution (5-10% normal goat serum or normal donkey serum). Sections were then incubated with primary antibodies against carbonic anhydrase 3 (1:150, SantaCruz), keratin 19 (1:50, DSHB, TROMA-III), COMP (1:200, Abcam, ab231977) COL I (1:100, Abcam, ab34710), COL II (1:400, Fitzgerald Industries, R-CR008), COL X (1:500, Abcam, ab58632), aggrecan (1:50, Millipore, AB1031) in blocking buffer at 4° C overnight. ARGxx (1:200, Abcam, ab3773) and CS (Abcam ab11570) used a MOM kit (Vector Laboratories, Burlingame, CA, USA) for staining and incubation of primary antibodies. Sections were washed with PBS and incubated with conjugated secondary antibody Alexa-Fluor-594 (Jackson ImmunoResearch Lab, Inc., 1:700) for 1 hr at room temperature. Sections were also stained to detect activity of tartrate-resistant acid phosphatase (1:80, Invitrogen, E6601A) and alkaline phosphatase (Vector Laboratories, SK- 5300). All sections were mounted with DAPI (Fisher Scientific, P36934) and imaged using Axio Imager 2 microscope equipped with AxioCam MRm monochrome camera with a 5x/0.15 N-Achroplan or 20x/0.5 EC Plan-Neofluar objectives and Zen2TM software (Carl Zeiss). For quantification, percent-stained area, cell count, and cell band area were calculated by thresholding, transforming the image to binary, and measuring particle number and area using ImageJ software, v1.53a, last access 9/11/2021 https://imagej.nih.gov/ij/.

### Tissue RNA isolation and microarray analysis

AF tissue from lumbar discs (L1-L6) of 7-month-old Wild-type (n=4 female) and ABCC6-/- mice (n=4 female) were collected and pooled for each individual mouse. RNA was isolated using RNeasy Mini Kit (Qiagen). Purified RNA was used to prepare cDNA using the GeneChip WT Plus kit (Thermo Fisher) and hybridized using Mouse Clariom S gene chips. Gene chips were scanned using an Affymetrix Gene Chip Scanner 3000 7G and Command Console Software. Experimental quality control, analyses and visualizations was performed using Transcriptome Analysis Console v4.0.2 (TAC). ABCC6-/- samples were compared to WT samples and included probesets where at least 50% of the samples had a DABG (detected above background) of p ≤ 0.05. Inclusion cutoffs were defined at a 1.75-fold change and p-value ≤ 0.1. Enrichment analysis of annotated genes was performed using PANTHER overrepresentation test, GO biological and molecular database annotations, and binomial statistical test with FDR ≤ 0.05. Array data are deposited in the GEO database (GSE188943).

### Biomechanical analysis

Lumbar vertebrae (L1-L2) from 23-week-old mice (*n=10 WT, 9 KO, 4 KO 2mM, 8 KO 40mM*) were isolated and stored in PBS-soaked gauze at -20° C before use. Samples were went through two freeze-thaw cycles and were μCT scanned before testing. Mechanical loading was conducted using a material testing system (TA Systems Electroforce 3200 Series II). First, each vertebrae were individually potted into a 2-mm plastic ring mold using acrylic resin (Ortho-Jet, Patterson Dental, Saint Paul, MN). Next, a 0.4-N compressive preload was applied, followed by a monotonic displacement ramp at 0.1 mm/s until failure. Force-displacement data was digitally captured at 25 Hz and converted to stress-strain using a custom GNU Octave script with microCT based geometric measurements, as previously described [72].

### Statistical analysis

All statistical analyses were conducted using Prism 8 (GraphPad). Graphical data are represented as mean ± SD, distribution was checked with Shapiro-Wilk test for normality, and differences between two groups were assessed by *t-*test or Mann-Whitney test when appropriate. Differences between the distribution of percentages was assessed using Chi-squared test, and the differences between more than 2 groups were assessed by ANOVA or Kruskal-Wallis followed by uncorrected Dunn’s multiple comparison test for non-normally distributed data. Statistical significance was considered p ≤ 0.05.

## Supporting Information

**S1 Fig. ABCC6^-/-^ mice exhibit trabecular thinning of caudal vertebrae.** Representative microCT reconstructions of *(a-h)* hemi- and *(a’-h’)* cross-sections show consistent trabecular thinning in caudal vertebrae of ABCC6^-/-^ mice at all timepoints. Quantitative microCT analysis of trabecular bone *(i-n)* BV/TV, Tb.Th., Tb.N., Tb.Sp., SMI, BMD, and cortical bone *(o-r)* B.Ar., MMi, Cs.Th., TMD. *(s)* Vertebral length, *(t)* disc height and *(u)* DHI are shown for caudal motion segments. Quantitative analyses are shown as mean ±SD (n = 2-4 discs/mice; 3-5 vertebrae/mice, n ≥ 5 mice/genotype). Significance was determined using unpaired t-test or Mann Whitney as appropriate. Scale bar=1 mm. BV/TV= bone volume/tissue volume. Tb.Th.= trabecular thickness. Tb.N.= trabecular number. Tb.Sp.= trabecular spacing. SMI= structural model index. BMD= bone mineral density. B.Ar.= bone area. MMI= mean polar moment of inertia. Cs.Th.= cross-sectional thickness. DHI= disc height index.

**S2 Fig. Presence of robust Alizarin Red staining within AF of 16-18M ABCC6^-/-^ disc.** *(a)* Representative Alizarin Red staining showing robust free calcium staining within the AF region of a 16-18M ABCC6^-/-^ mouse disc.

**S3 Fig. Caudal discs of ABCC6^-/-^ mice showed minor AF degeneration with comparable collagen fiber composition.** *(a-b’)* Safranin O/Fast Green staining of *(a-a’)* 7-8 month and *(b-b’)* 16–18-month-old lumbar discs showed tissue morphology and proteoglycan content consistent with age-related disc degeneration (row 1, scale bar = 200μm and rows 2-3, scale bar = 50μm). *(c, d)* Histological grading analysis using the modified Thompson scale showed altered levels of NP and AF degradation but similar average Thompson grades. *(e-g)* Picrosirius Red staining and quantification of percent collagen fiber area showed no difference in overall collagen fiber thickness in the AF of ABCC6^-/-^ caudal discs. Quantitative analyses are shown as mean ±SD. Significance of grading distribution was determined using a *x*^2^ test. Significance of average grade data and percent area were determined using unpaired t-test or Mann Whitney test as appropriate.

